# High quality genome assemblies of African cattle breeds using PacBio HiFi sequencing

**DOI:** 10.1101/2025.04.17.649430

**Authors:** Isidore Houaga, Meenu Bhati, Zabron Nziku, Athumani Nguluma, Ntanganedzeni Mapholi, Lucky T. Nesengani, Moses Ogugo, Duhamel C.Y. Sagbo, Loukaïya Zorobouragui, Mariano B.Y. Boco, Appolinaire Djikeng, James G.D. Prendergast, Gregor Gorjanc, Hannes Becher

**Affiliations:** The Roslin Institute, Royal (Dick) School of Veterinary Studies, University of Edinburgh, Easter Bush Campus, Midlothian, EH25 9RG, UK; Centre for Tropical Livestock Genetics and Health, Easter Bush, Midlothian, EH25 9RG, UK; Department of Agriculture and Animal Health, College of Agriculture and Environmental Sciences, The University of South Africa, Cnr Justice Mahomed & Steve Biko Streets, PO Box 392, Pretoria, South Africa; Tanzania Livestock Research Institute (TALIRI) Eastern Zone, Tanga, Tanzania; Department of Animal, Aquaculture, and Range Sciences, Sokoine University of Agriculture, Morogoro, Tanzania; The International Livestock Research Institute, PO Box 30709, Nairobi, Kenya; African Institute of Data Driven Animal and Plant Breeding and Genetics (DATAGENES-AFRICA), Aplahoue, Benin Republic; Laboratory of Ecology, Health and Animal Production (LESPA), Faculty of Agronomy, University of Parakou, Benin; Ferme d’Élevage de Samiondji BP: 50 Covè, Benin Republic

## Abstract

Africa has a uniquely rich cattle diversity of ∼150 breeds comprising the *Bos taurus indicus* sub-species, *Bos taurus taurus*, and their crosses. These represent ∼23% of the global cattle population. However, high quality, representative assemblies are limited for African cattle and especially for indicine breeds. Here we built high quality *de novo* assemblies for five important African indigenous cattle breeds using PacBio HiFi sequencing: Lagune (*Bos taurus taurus)*, Gudali, Iringa Red and Singida White (*Bos taurus indicus*), and Mpwapwa (*Bos taurus taurus x Bos taurus indicus*). These new assemblies are the most contiguous and complete African cattle assemblies produced so far, with genome sizes of 3.25 - 3.36Gb, contiguity N50s ranging from 83.59Mb to 97.87Mb and scaffold N50s from 100.30Mb to 113.37Mb. BUSCO genome completeness scores were also higher than 99.68%, indicative of highly contiguous assemblies. These improved and highly contiguous genome assemblies are consequently a valuable resource for future African and global livestock genomic studies.

## Background & Summary

Cattle play important roles globally and in Africa. Providing meat and milk, they are a significant source of nutrition and livelihood to many African societies. They are also used in agriculture as a source of traction and manure. Further, cattle are of great societal importance, including during birth, marriage, death, or traditional ceremonies, and are a representation of prestige and power^1,2^. Since their domestication around 10,000 years ago, hundreds of distinct cattle breeds have been developed, characterised by high variation in physical appearance, productivity levels, environmental adaptation and disease tolerance^3^. Africa has a uniquely rich cattle diversity with about 150 known cattle breeds or populations comprised of *Bos taurus taurus, Bos taurus indicus*, and their crosses^4^. These account for up to 23% of the global cattle population^5^, compared to only 8% for European cattle.

Despite the astounding diversity of African cattle, there is a conspicuous lack of high quality and representative assemblies, particularly for *Bos taurus indicus* breeds. Multiple introductions and migrations of both taurine and indicine cattle into Africa, combined with diverse environments, pathogens, and cultures have resulted in high genetic diversity among its cattle breeds^6,7^. However, this diversity is inadequately represented by currently published reference genomes, which comprise the Asian-derived breed Brahman^8^ and the European breeds Angus^8^ and Hereford^9^. A recent study generated the first African cattle genome assemblies for the West African Taurine N’Dama and an East African stabilized cross between indicine and taurine breeds, called Ankole. This study identified an extra 116.1 Mb (4.2%) of sequence absent from the current Hereford reference sequence^9^ and consequently inaccessible in previous studies^6^. However, the new assemblies for N’Dama and Ankole had modest contiguities, N50 of 10.7-18.6 Mb. Gene content completeness was 92.6% - 93.1%, using Benchmarking Universal Single-Copy Orthologs (BUSCO), suggesting that more high quality and representative assemblies are needed for African cattle breeds. Such high quality and representative assemblies are expected to improve read mapping, allelic biases, structural variant calling, as well as downstream genomic analyses such as detecting signatures of selection^6,10^.

In this study, we address the current lack of high-quality reference genomes for African cattle breeds. Using PacBio HiFi sequencing data, we generated state-of-the-art *de novo* genome assemblies for five important African indigenous cattle breeds. These are Lagune (*Bos taurus taurus*), Gudali, Iringa Red and Singida White (all three *Bos taurus indicus*), and Mpwapwa (*Bos taurus taurus x Bos taurus indicus*). The five breeds are shown in Fig. 1. The Lagune and Gudali cattle breeds were sampled in the Republic of Benin, West Africa, while Iringa Red, Singida White and Mpwapwa cattle breeds were sampled in Tanzania, East Africa. **Lagune** is a small hardy breed known for its adaptability to humid tropical climates and resistance to diseases such as trypanosomiasis making it valuable for meat production in tsetse fly-infested regions^11^. **Gudali** is a robust breed originating from West and Central Africa, known as an excellent dairy producer. It is also known for its strength and adaptability to semi-arid regions and is used for draft purposes^12^. **Iringa Red** is native to Tanzania, prized for its drought resistance and ability to thrive in challenging environments, making it useful for meat production and agricultural work^13^. **Singida White** is a Tanzanian indigenous breed well adapted to harsh environments, known for its use in meat production and as farmer-preferred draft animals in rural areas^14^. **Mpwapwa** is a dual-purpose (milk and beef) breed developed in Tanzania, bred for resilience in tropical climates and resistance to common diseases, enhancing food security in rural communities^13^. The Mpwapwa, Iringa Red and Singida White breeds have never been subject to any whole-genome sequencing study in Tanzania.

**Fig. 1.**
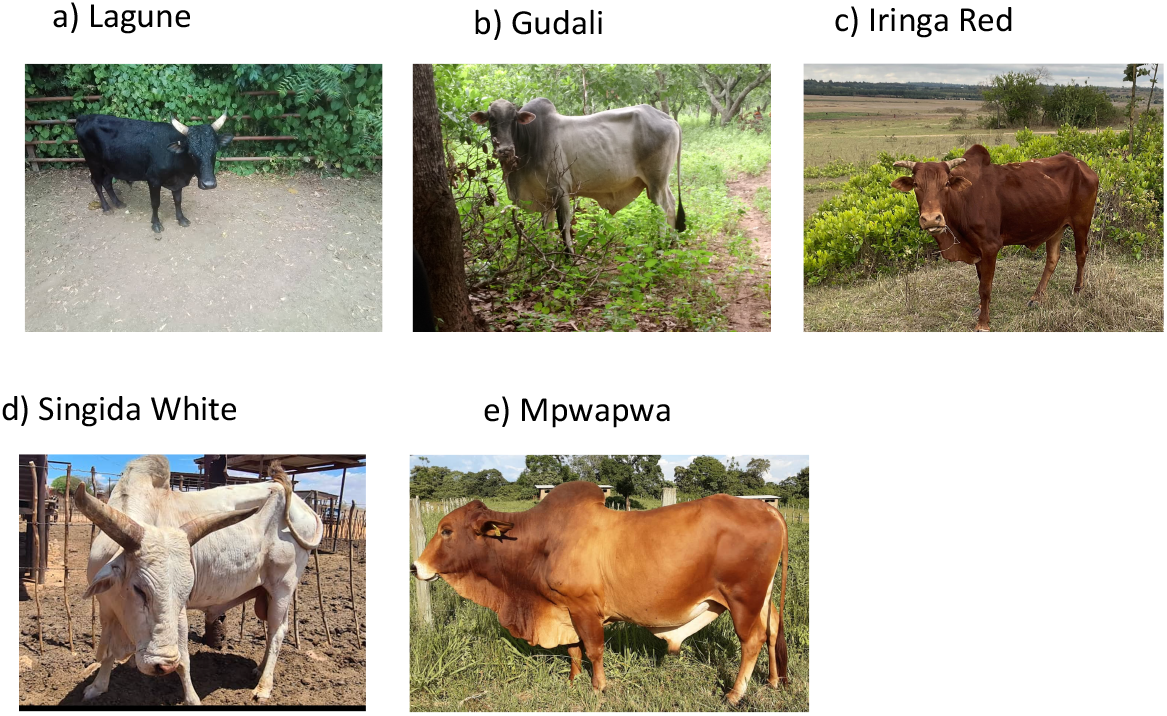
Pictures of the five sequenced cattle breeds: a) Lagune (Bull 511), b) Gudali, c) Iringa Red, d) Singida White and e) Mpwapwa (Bull NAIC 906).

Our new haplotype-resolved assemblies are the most contiguous and complete African cattle assemblies produced so far with genome size ranging from 3.25 to 3.36 Gb and with heterozygosity ranging between 0.37% (Lagune) and 0.68% (Iringa Red). Contiguity N50 ranged from 83.59 Mb (Iringa Red) to 97.87 Mb (Lagune) and scaffold N50 from 100.30 Mb (Iringa Red) to 113.37Mb (Lagune). BUSCO genome completeness as estimated by *Compleasm* using the mammalian set (n=9,226) was higher than 99.68% for all assemblies. Assembly accuracies, assessed with Merqury, exceeded QV 67 (99.9999%). Across the five indigenous cattle genomes, an average of 1.84 Gb of repeat sequences were identified, representing approximately 56% of each genome, with retroelements being the most abundant repeat class. These improved and highly accurate genome assemblies will provide valuable resources for African and global genomic studies of cattle and livestock in general.

## Methods

### Ethics statement

The study was approved by the Tanzania Livestock Research Institute Ethical Clearance Committee (Reference number: TLRI/CC.21/024). The blood collection was carried out by trained local veterinarians, according to the approved institutional guidelines. We obtained consent from all farmers before collecting blood samples.

### Animal sampling

In this study, a Lagune bull (ID:511, born on 20/01/2020) was selected at the State-owned Samiondji Breeding Farm (07°26’30” N, 02°22’59” E) in Zagnanado Commune, in the Zou Department, southern-central Benin. A Gudali cow was selected from a smallholder farmer at Kasuala (8°53’18.96’’N 2°44’56.796’’E), Commune of Tchaourou, in the Borgou Department, northern-east Benin. The Iringa Red bull was selected from a smallholder farmer at Ibumila (8°3’28.17” S, 35° 33’47.65” E), Iringa Region, Central Tanzania. The Singida White cattle breed cow (ID:01) was selected at the Tanzania Livestock Research Institute, research station located in Kongwa (6°12’0” S, 36°25’0” E), Dodoma region, Tanzania. A Mpwapwa cattle breed bull (NAIC 906, born on 26/06/2013) was selected at the Tanzania National Artificial Insemination Centre (3°22’12” S, 36°50’09” E) located in the Usa River, Arusha, Northen Tanzania. Blood samples were collected from the jugular vein into a labelled 5 ml EDTA vacutainer tubes and immediately transferred to the laboratory in a cool box containing icepack and stored at -20°C until DNA extraction.

### DNA extraction and PacBio long-reads Sequencing

The blood samples were allowed to thaw for 30 minutes at room temperature. High Molecular Weight (HMW) DNA was extracted from 200 μL bovine whole blood using the Pacific Biosciences (PacBio) HMW DNA extraction from mammalian cells – Nanobind® tissue kit (102-302-100) following manufacturer’s instructions. The PacBio SMRTBell® prep kit 3.0 was used for library preparation and the Revio polymerase kit was used for loading the libraries onto the sequencer. The samples were sequenced at 225 pM on the PacBio Revio platform at Edinburgh Genomics using two Revio SMRT Cells per sample. The sequencing yielded a total of 8.2-12.6 M reads and 160-219 Gbp, corresponding to a genomic coverage depth of approximately 53-72X. The summary statistics of the sequencing reads are presented in Table 1.

**Table 1.** Summary statistics of the raw sequencing data.

| Cattle breed | Subpecies | Country/Region | Sex | HiFi Yield (Gb) | Mean HiFi RL (bp) | Median QV | Coverage |
| --- | --- | --- | --- | --- | --- | --- | --- |
| Lagune | *Bos taurus* | Benin/We st-Africa | ♂ | 218 | 19,476 | 32 | 72x |
| Gudali | *Bos indicus* | Benin/We st-Africa | ♀ | 219 | 19,905 | 34 | 73x |
| Iringa Red | *Bos indicus* | Tanzania/ East-Africa | ♂ | 160 | 19,479 | 32 | 53x |
| Singida White | *Bos indicus* | Tanzania/ East-Africa | ♀ | 190 | 15,147 | 33 | 63x |
| Mpwapwa | *Bos taurus* x *Bos indicus* | Tanzania/ East-Africa | ♂ | 204 | 18,195 | 33 | 68x |

RL: Read Length.

### Genome assembly

De novo assembly was run with the 53-72X coverage obtained from each sample using hifiasm (v0.19.9)^15^ with default parameter except for trimming the first 20 base pairs. The assemblies’ contigs were scaffolded using the recently published Hereford reference cattle genome (ARS-UCD2.0) using RagTag (v 2.1.0)^16^ without the correction step because of the high quality of our assemblies. Contaminant contigs identified and flagged during the NCBI submission process were removed from the final genome assemblies. These contigs were all unplaced and did not correspond to autosomal, sex chromosomal, or mitochondrial sequences, ensuring that only high-confidence genomic regions were retained.

### Repeat sequences and annotation

Repeat annotation of the five African cattle genome assemblies was performed using RepeatMasker (v4.1.5)^17^ with established repeat libraries, including Dfam (v3.8)^18^ and the RepBase RepeatMasker Edition (20181026) and tandem repeats with TRF (v4.09.1). De novo repeat identification from RepeatModeler (v2.0.5)^19^ was not integrated due to the high coverage provided by these curated libraries, resulting in only 0.01% of unidentified repeats.

## Data Records

This whole genome sequencing project has been deposited at DDBJ/ENA/GenBank under the accession XXXXXX000000000. The version described in this paper is version XXXXXX010000000. The five scaffolded primary genome assemblies are accessible under BioProject ID PRJNA1197437. The Lagune, Gudali, Iringa Red, Singida White and Mpwapwa genome assemblies have been respectively assigned accessions SAMN45804670, SAMN45804671, SAMN45804673, SAMN45804674 and SAMN45804672. The raw PacBio Hifi sequencing reads will be submitted to the Sequence Read Archive (SRA) database.

## Technical Validation

### African cattle genome assemblies and quality assessment

We used FastQC (v0.12.1, https://www.bioinformatics.babraham.ac.uk/projects/fastqc/) to assess the quality of the sequencing reads using default parameters, and summarised the results using multiQC^20^. This did not result in any evidence for sequencing adapter contamination. After assembling the genomes, we employed the GenomeScope (v 2.0)^21^ for genome profiling using 21-mer counts generated by Meryl (v1.2; https://github.com/marbl/meryl) from the HiFi sequencing reads, see Fig. 2. We evaluated the quality of the primary genome assembly using Merqury (v1.1)^22^, see Fig. 3. The spectra are used to compare k-mers from the sequencing reads with those found in the haplotype-resolved genome assemblies, providing insights into assembly completeness, missing data, and errors. Across all breeds, the k-mer spetra show two prominent peaks as expected for diploids individuals with some degree of heterozygosity. In each plot, shared k-mers in green dominate in the right-hand diploid 2x peak, whereas haplotype-specific k-mers shown in red and blue dominate the left-hand haploid 1x peak. K-mers present in the sequencing data but absent from the final assemblies are shown in grey. These are largely of low multiplicity, at the very left of each plot, indicating that sequencing errors and contamination were not erroneously incorporated and that the assemblies are highly complete. The completeness and integrity of the genome assemblies were further evaluated using the BUSCO implemented in compleasm^23^ using the mammalian dataset “mammalia_odb10” (n=9226). The evaluation found that 99.68% (9195) to 99.93 % (9220) of the core mammalian genes were present in the African cattle genomes, including 98.46 to 98.65 % single-copy, 1.19% to 1.47% duplicated, 0.03% fragmented and 0.03% to 0.29 % missing genes from the mammalian data set (Table 2). The k-mer databases were used to calculate the overall quality values of the assemblies using Merqury^18^. The assemblies had high accuracy with QV 67.5-QV 71.7 (>99.9999%, Table 2). These confirmed the completeness of the five African cattle breeds reported in this manuscript. The QUAST^24^ software (v5.2.0) calculated that the five new assemblies cover 95.2% (GUD and SIW), 95.6% (IRR), 95.8% (MPW) and 96.4% (LAG) of the ARS-UCD2.0 (the most recent Hereford genome) higher than the reported 93.9% and 94.0% of the ARS-UCD1.2 Hereford genome covered respectively by the African N’Dama and Ankole cattle breeds^6^. The genome lengths ranged from 3.25 Gb (GUD) to 3.36 Gb (SIW) with 340 (MPW) to 622 (SIW) contigs. The contig N50 ranged from 83.59 Mb (Iringa Red) to 97.87 Mb (Lagune) higher than 10.70 Mb and 18.60 Mb previously reported for the N’Dama and the Ankole cattle breeds respectively^6^. The scaffolded assemblies had 222 (GUD) to 446 (SIW) scaffolds in total with scaffold N50 of 100.30 Mb (Iringa Red) to 113.37 Mb (Lagune) as shown in Table 2. The scaffold N50 of Lagune (113.37), Singida White (108.17) and Mpwapwa (105.95 Mb) are higher than the values of 104.80 Mb and 84.50 Mb reported for the N’Dama and Ankole genomes, respectively^6^. These statistics and the above quality assessment metrics confirm that our new assemblies are the most contiguous and complete African cattle assemblies produced so far.

**Table 2.** Genome assembly statistics of five African cattle breeds.

| Statistics | Lagune | Gudali | Iringa Red | Singida White | Mpwapwa |
| --- | --- | --- | --- | --- | --- |
| Raw primary assemblies as from *hifiasm*<sup>15</sup> |  |  |  |  |  |
| Consensus accuracy | QV67.5 | QV69.6 | QV68.7 | QV68.9 | QV71.7 |
| Genome size (bp) | 3,338,004,939 | 3,252,386,270 | 3,302,534,872 | 3,356,428,914 | 3,296,393,950 |
| GC content (%) | 44.01 | 43.95 | 44.09 | 44.26 | 44.09 |
| Heterozygosity | 0.37 | 0.46 | 0.68 | 0.47 | 0.66 |
| #Contigs | 456 | 392 | 414 | 622 | 340 |
| Average contig length (bp) | 7,320,186 | 8,296,903 | 7,977,137 | 5,396,187 | 9695276 |
| Contig N50 | 97,870,685 | 94,506,920 | 83,585,658 | 91,475,713 | 96,250,906 |
| Contig L50 | 14 | 14 | 15 | 15 | 14 |
| Largest contig | 173,713,701 | 173,964,285 | 161,906,315 | 172,444,127 | 164,144,583 |
| Smallest contig | 15,073 | 16,340 | 5,341 | 11,215 | 16,271 |
| **Clean scaffolded primary genome assemblies deposited at GenBank** |  |  |  |  |  |
| #Scaffolds | 292 | 222 | 308 | 446 | 302 |
| Total scaffold length (bp) | 3,327,642,104 | 3,239,885,270 | 3,296,547,564 | 3,345,996,902 | 3,295,703,406 |
| Average scaffold length (bp) | 11,396,034.6 | 14,594,077.79 | 10,703,076.51 | 7,502,235.21 | 10,912,925.19 |
| scaffold N50 | 113,365,931 | 100,524,608 | 100,300,525 | 108,170,087 | 105,951,773 |
| scaffold L50 | 13 | 13 | 13 | 13 | 13 |
| Largest scaffold | 173,713,701 | 173,964,285 | 161,906,315 | 172,444,127 | 166,841,832 |
| Smallest scaffold | 15,073 | 16,340 | 20,870 | 11,215 | 16,271 |
| ARS-UCD2.0 Genome fraction (%) | 96.4 | 95.2 | 95.6 | 95.2 | 95.8 |
| # Completeness (Compleasm), library: mammalia_odb10 (n=9,226) | 9,220 (99.93%) | 9,220 (99.94%) | 9,195 (99.68%) | 9,220 (99.93%) | 9,220 (99.93%) |
| Complete and Single copy | 9,100 (98.63%) | 9,101 (98.65%) | 9,085 (98.47%) | 9,084 (98.46%) | 9,097 (98.60%) |
| Complete and duplicated | 120 (1.3%) | 119 (1.29%) | 110 (1.19%) | 136 (1.47%) | 123 (1.33%) |
| Fragmented | 3 (0.03%) | 3 (0.03%) | 4 (0.04%) | 3 (0.03%) | 3 (0.03%) |
| Missing | 3 (0.03%) | 3 (0.03%) | 27 (0.29%) | 3 (0.03%) | 3 (0.03%) |

**Fig. 2.**
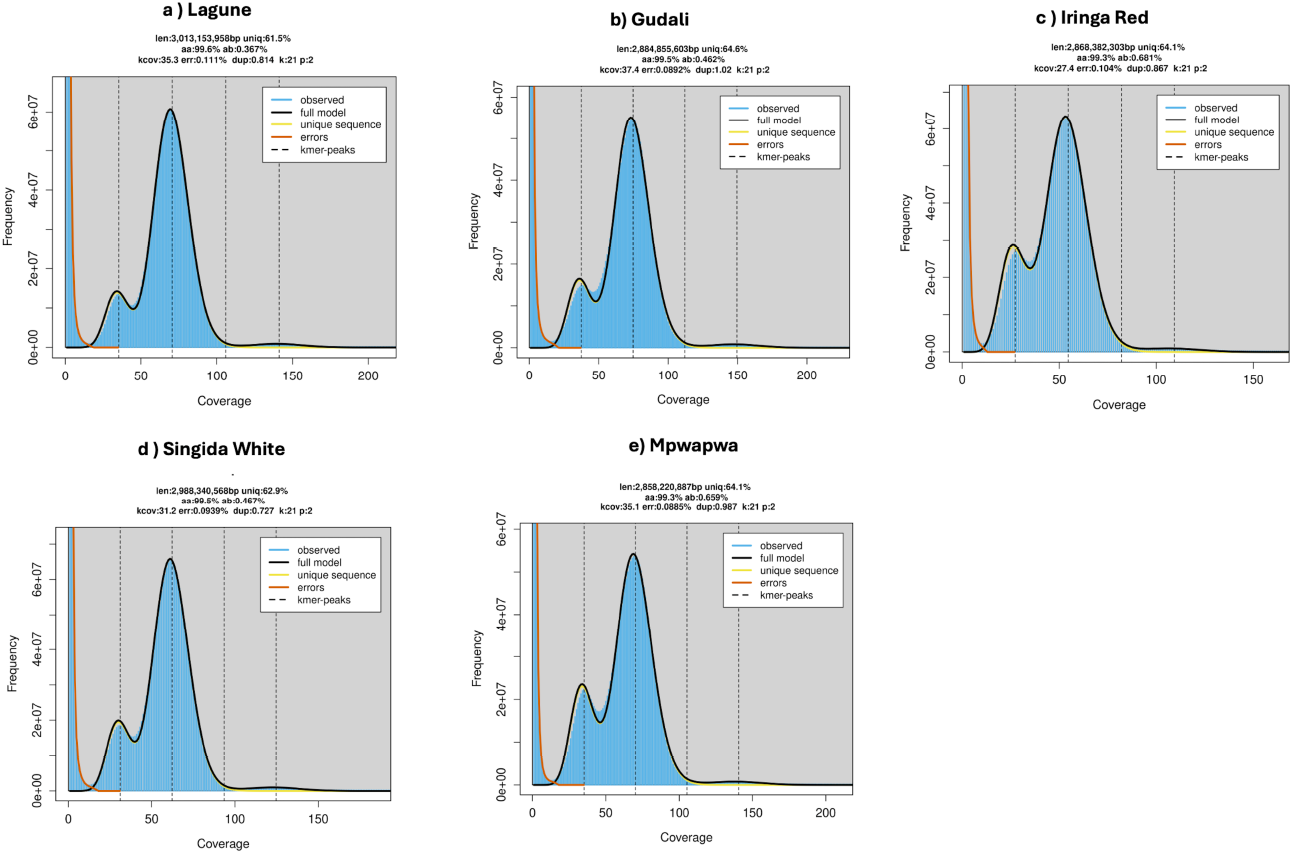
GenomeScope profiles for five novel African cattle genome assemblies. The estimated genome size ranged between 2.88 (Gudali) and 3.01 Gb (Lagune) with heterozygosity between 0.36 (Lagune) and 0.68% (Iringa Red).

**Fig. 3.**
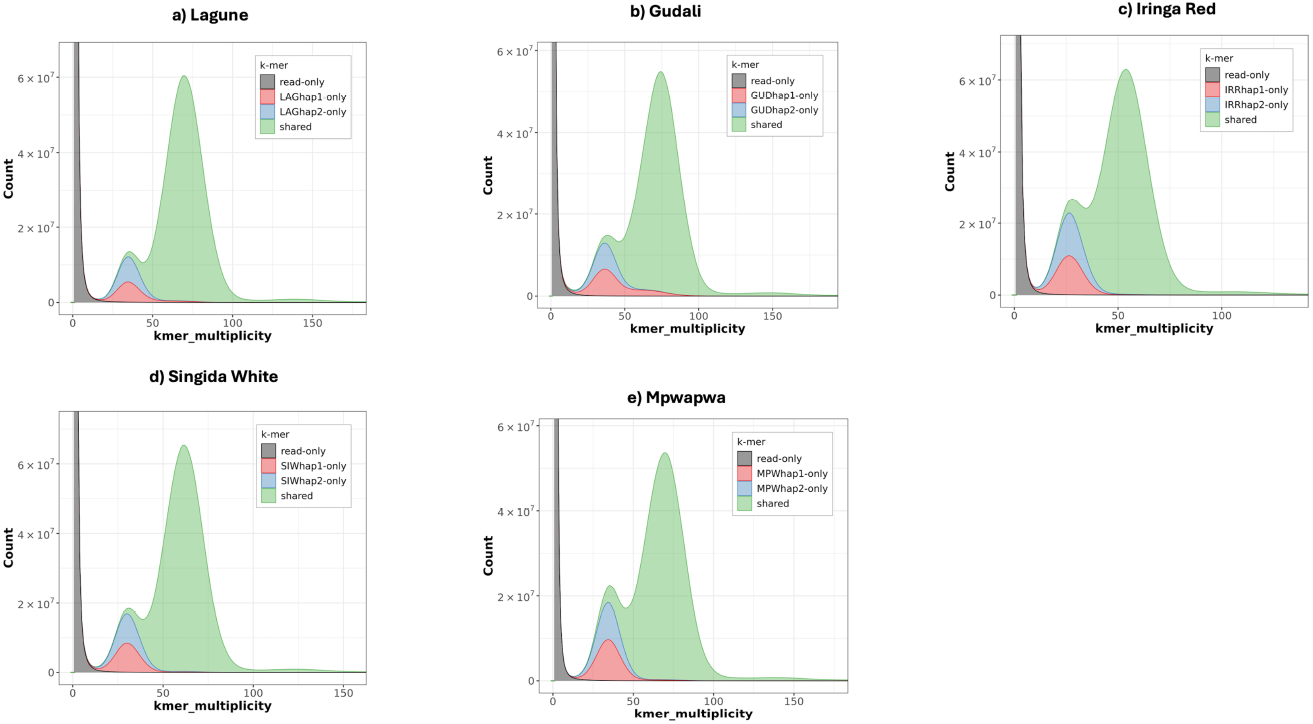
Merqury k-mer spectrum copy number (CN) plots for five haplotype-resolved African cattle genome assemblies. K-mers shared between haplogenomes are shown in green, those private to one haplogenome are shown in blue and red. K-mers absent from the assembly are shown in grey.

### Genomic relationships

We generated a dendrogram by comparing the assemblies’ k-mer contents to other publicly available genomes from NCBI using mashtree^25^. The publicly available assemblies include outgroups (Human-GCA000001405.29 GRCh38.p14, Goat-GCA 001704415.2 ARS1.2, Sheep-ARS-UI Ramb v3.0, African Buffalo-GCA 902825105.1 and Water buffalo-GCF 019923935. 1 NDDB SH 1); *Bos indicus* (GCA_026262515.1 ASM2626251v1, GCA_026262465.1 ASM2626246v1, GCA 030269505. 1 ASM3026950v1, GCA 003369695.2 Brahman 1 and GCA 000247795.2 1.0); *Bos indicus × Bos taurus* (Ankole GCA_905123885) and *Bos taurus* (NDama GCA_905123515, Jersey GCA_021234555.1 ARS, GCA 021347905.1 ARS Holstein-Friesian 1, GCA 003369685.2 UOA Angus 1, Hereford ARS-UCD2.0, GCA 028973685.2 SNU Hanwoo 2.0, *GCA 040286185*.*1 ASM4028618v1* and GCA 018282465.1 ARS Simm1.0).

The genome-wide similarities between the new African cattle assemblies (blue) and publicly available assemblies are shown in Fig. 4. The new African taurine assembly (Lagune) was closely related to the African N’Dama taurine. The new assemblies of indicine origin and Bos indicus x Bos taurus crossbred also clustered together.

**Fig. 4.**
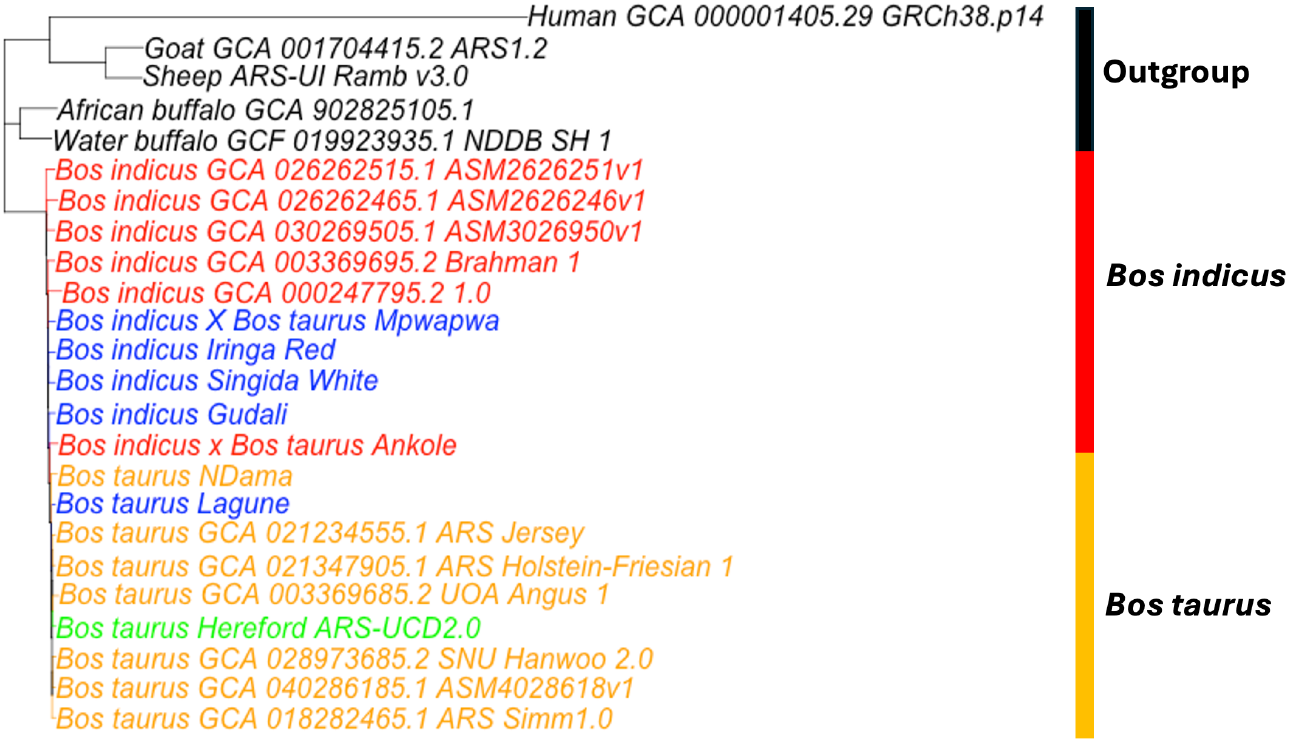
Genome similarities between new African cattle assemblies and publicly available assemblies. The new African assemblies generated here are in blue.

### Repeat sequences and annotation

The repeat content across the five African cattle genome assemblies is presented in Table 3. The total proportion of masked bases, representing repetitive DNA, ranged from 55.7% to 56.9%, with Singida White showing the highest proportion of repeat content (56.9%), and Gudali the lowest (55.7%). The repeat contents are slightly higher than the repeat content reported for previously characterized bovine genomes such as *Bos taurus*, which is approximately 47%^8^. Retroelements represented the largest class of repeats, contributing approximately 38–39% of the genome across all breeds. Among retroelements, LINEs (Long Interspersed Nuclear Elements) were the most abundant subtype, accounting for about 23–23.5%, followed by SINEs (Short Interspersed Nuclear Elements) at 9.6–9.9%, and LTR elements (Long Terminal Repeats) at 5.4–5.5%. Satellite DNA, a key component of centromeric and pericentromeric regions, displayed greater variability, ranging from 13.9% in Gudali to 16.2% in Singida White. Overall, the Mpwapwa breed had the highest repeat content in absolute terms, while Gudali had the lowest. Despite these differences, the general repeat composition and distribution patterns were broadly similar across all five cattle genomes, reflecting conserved repeat architecture among these breeds.

**Table 3.** Repeat sequences contents across five African cattle breeds. Percentages are calculated based on the total genome length of each assembly, and small summation discrepancies may occur due to overlapping annotations, or repeats classified in multiple categories by Repeatmasker.

| Main | subtype | Lagune | Gudali | Iringa Red | Singida White | Mpwapwa |
| --- | --- | --- | --- | --- | --- | --- |
| Retroelements |  | 1,286,930,678<br>(38.7%) | 1,260,581,845<br>(38.9%) | 1,266,327,947<br>(38.4%) | 1,235,053,136<br>(37.9%) | 1,275,256,348<br>(38.7%) |
|  | LINEs | 777,966,277<br>(23.4%) | 761,124,491<br>(23.5%) | 765,251,253<br>(23.2%) | 746,988,345<br>(22.9%) | 775,394,625<br>(23.5%) |
|  | LTR | 184,078,941<br>(5.5%) | 178,101,720<br>(5.5%) | 179,680,343<br>(5.5%) | 174,600,597<br>(5.4%) | 176,345,576<br>(5.4%) |
|  | PLEs | 57,571 (0.0%) | 55,168 (0.0%) | 54,606 (0.0%) | 53,590 (0.0%) | 54,058 (0.0%) |
|  | SINEs | 324,885,460<br>(9.8%) | 321,355,634<br>(9.9%) | 321,396,351<br>(9.7%) | 313,464,194<br>(9.6%) | 323,516,147<br>(9.8%) |
| DNA Transposons |  | 61,865,922<br>(1.9%) | 61,564,772<br>(1.9%) | 61,401,094<br>(1.9%) | 59,996,704<br>(1.8%) | 61,763,643<br>(1.9%) |
| Rolling Circles |  | 290,862<br>(0.0%) | 288,143<br>(0.0%) | 292,224 (0.0%) | 282,605<br>(0.0%) | 289,875<br>(0.0%) |
| Unclassified |  | 525,581<br>(0.0%) | 479,986<br>(0.0%) | 530,409 (0.0%) | 468,391<br>(0.0%) | 544,117<br>(0.0%) |
| Total Interspersed Repeats |  | 1,349,379,752<br>(40.6%) | 1,322,681,771<br>(40.8%) | 1,328,314,056<br>(40.3%) | 1,295,571,821<br>(39.7%) | 1,337,618,166<br>(40.6%) |
| Satellites |  | 475,498,670<br>(14.3%) | 451,417,569<br>(13.9%) | 491,232,699<br>(14.9%) | 528,409,710<br>(16.2%) | 467,096,074<br>(14.2%) |
| Simple Repeats |  | 27,545,432<br>(0.8%) | 25,013,594<br>(0.8%) | 26,856,901<br>(0.8%) | 24,583,321<br>(0.8%) | 27,251,561<br>(0.8%) |
| Low Complexity |  | 4,497,462<br>(0.1%) | 4,273,074<br>(0.1%) | 4,333,715 (0.1%) | 4,256,759<br>(0.1%) | 4,358,856<br>(0.1%) |
| Small RNA |  | 45,297,364<br>(1.4%) | 44,731,650<br>(1.4%) | 44,678,161<br>(1.4%) | 43,653,726<br>(1.3%) | 45,209,784<br>(1.4%) |
| Total Repeats |  | 1,858,696,960<br>(55.9%) | 1,804,973,058<br>(55.7%) | 1,852,269,687<br>(56.2%) | 1,854,826,538<br>(56.9%) | 1,838,123,418<br>(55.8%) |

**LINEs:** Long Interspersed Nuclear Elements; **SINEs:** Short Interspersed Nuclear Elements; **LTR:** Long Terminal Repeats, **PLEs:** Penelope-like Elements.

## Code Availability

Parameters for all the commands used to assemble and analyse the genomes were the default parameters of the corresponding software, unless mentioned and where necessary.

## Acknowledgements

The authors acknowledge funding from the Royal Society Newton International Fellowship (NIF/R1/201150) and the Roslin Foundation grant (RF2303) to IH, the Global Challenge Research Fund-Official Development Assistance (GCRF-ODA) to AD, the Bill & Melinda Gates Foundation and UK aid from the UK Foreign, Commonwealth and Development Office (INV-040641) under the auspices of the Centre for Tropical Livestock Genetics and Health (CTLGH), established jointly by the University of Edinburgh, Scotland’s Rural College (SRUC), and the International Livestock Research Institute. The findings and conclusions contained within are those of the authors and do not necessarily reflect positions or policies of the Bill & Melinda Gates Foundation nor the UK Government. Sequencing was carried out by Edinburgh Genomics, The University of Edinburgh, which is partly supported with core funding from the Natural Environment Research Council (UKSBS PR18037). MB, JP, GG, and HB acknowledge support from BBSRC Institute Strategic Programme funding to the Roslin Institute (BBS/E/D/30002275, BBS/E/D/10002070, BBS/E/RL/230001A, BBS/E/RL/230001B, and BBS/E/RL/230001C) and the University of Edinburgh. MB is supported by the Swiss National Science Foundation (SNSF) as a Postdoc Mobility fellow (grant no. 222130).

## Author contributions

IH, MB, JP, GG and HB designed the study. IH assembled the genomes, conducted data analysis and wrote the first draft of the manuscript. MB contributed to genome assembly, data analysis and conducted repeat content analysis. ZN, AN, DCYS, LZ, MBYB collected samples. MO extracted high molecular weight DNA. NM and LTN advised on data analysis steps. All authors reviewed the manuscript.

## References

1. di Lernia, S. et al. Inside the “African cattle complex”: Animal burials in the Holocene central Sahara. PLoS One 8, e56879 (2013).

2. Schneider, H.K. The subsistence role of cattle among the Pakot and in East Africa. American Anthropologist. 59, 278–300 (1957).

3. Ajmone-Marsan, P., Lenstra, J.A. & Fernando Garcia, J. On the origin of cattle: how aurochs became domestic and colonized the world. Evolutionary Anthropologist 19, 148–157 (2010).

4. Mwai, O., Hanotte, O.Kwon, Y.-J. & Cho, S. African indigenous cattle: unique genetic resources in a rapidly changing world. Asian-Australasian Journal of Animal Science 28, 911–921 (2015).

5. De Boer, H. Cattle genetic resources. Livestock Production Science 29, 256–258 (1991).

6. Talenti, A., Powell, J., Hemmink, J.D. et al. A cattle graph genome incorporating global breed diversity. Nature Communications 13, 910 (2022).

7. Kim, K., Kwon, T., Dessie, T. et al. The mosaic genome of indigenous African cattle as a unique genetic resource for African pastoralism. Nature Genetics 52, 1099–1110 (2020).

8. Koren, S., Rhie, A., Walenz, B. et al. De novo assembly of haplotype-resolved genomes with trio binning. Nature Biotechnology 36, 1174–1182 (2018).

9. Rosen, B. D. et al. De novo assembly of the cattle reference genome with single-molecule sequencing. GigaScience 9, 1–9 (2020).

10. Lloret-Villas, A., Bhati, M., Kadri, N.K. et al. Investigating the impact of reference assembly choice on genomic analyses in a cattle breed. BMC Genomics 22, 363 (2021).

11. Ahozonlin, M. C., Gbangboche, A. B., & Dossa, L. H. Current Knowledge on the Lagune Cattle Breed in Benin: A State of the Art Review. Ruminants 2, 271–281.

12. Zorobouragui, L., Houaga, I., Assani, A.S., Worogo, H.S.S., Kinkpe, L., Periasamy, K. & Alkoiret, I.T. Breeding practices and selection criteria in Gudali cattle breed from Benin: implications for the design of a community-based breeding program. Frontiers in Animal Science 5,1454071 (2025).

13. Wilson, R.T. When is a “breed” not a breed: the myth of the Mpwapwa cattle of Tanzania. Tropical Animal Health and Production 53, 233 (2021).

14. Salum, K. A., Laswai, G. H. & Mushi, D. E. Performance of Boran and Two Strains of Tanzania Short Horn Zebu Cattle Fed on Three Different Diets. International Journal of Animal Science and Technology 8, 21–29 (2024).

15. Cheng, H., Concepcion, G.T., Feng, X., Zhang, H. & Li, H. Haplotype-resolved de novo assembly using phased assembly graphs with hifiasm. Nature Methods 18,170-175 (2021).

16. Alonge, M. et al. Automated assembly scaffolding using RagTag elevates a new tomato system for high-throughput genome editing. Genome Biology 23, 258(2022).

17. Chen, N. Using Repeat Masker to identify repetitive elements in genomic sequences. Current protocols in bioinformatics 5, 4.10. 11–14.10. 14 (2004).

18. Wheeler, T.J. et al. Dfam: a database of repetitive DNA based on profile hidden Markov models. Nucleic Acids Research 41, D70–D82 (2013).

19. Flynn, J.M. et al. RepeatModeler2 for automated genomic discovery of transposable element families. Proceedings of the National Academy of Sciences 117, 9451–9457.

20. Ewels, P., Magnusson, M., Lundin, S., & Käller, M. MultiQC: Summarize analysis results for multiple tools and samples in a single report. Bioinformatics, 32, 3047–3048 (2016).

21. Ranallo-Benavidez, T.R., Jaron, K.S. & Schatz, M.C. GenomeScope 2.0 and Smudgeplot for reference-free profiling of polyploid genomes. Nature Communications 11, 1432 (2020).

22. Rhie, A., Walenz, B. P., Koren, S. & Phillippy, A. M. Merqury: reference-free quality, completeness, and phasing assessment for genome assemblies. Genome Biology 21, 245 (2020).

23. Neng Huang, Heng Li, compleasm: a faster and more accurate reimplementation of BUSCO. Bioinformatics 39, btad595 (2023).

24. Mikheenko, A., Prjibelski, A., Saveliev, V., Antipov, D. & Gurevich, A. Versatile genome assembly evaluation with QUAST-LG. Bioinformatics 34, i142–i150 (2018).

25. Katz, L. S. et al. Mashtree: a rapid comparison of whole genome sequence files. Journal of Open-Source Software 4, 1762 (2019).

